# PIKfyve inhibition blocks endolysosomal escape of α-synuclein fibrils and spread of α-synuclein aggregation

**DOI:** 10.1101/2021.01.21.427704

**Authors:** Stephanie K. See, Merissa Chen, Sophie Bax, Ruilin Tian, Amanda Woerman, Eric Tse, Isabel E. Johnson, Carlos Nowotny, Elise N. Muñoz, Janine Sengstack, Daniel R. Southworth, Jason E. Gestwicki, Manuel D. Leonetti, Martin Kampmann

## Abstract

The inter-cellular prion-like propagation of α-synuclein aggregation is emerging as an important mechanism driving the progression of neurodegenerative diseases including Parkinson’s disease and multiple system atrophy (MSA). To discover therapeutic strategies reducing the spread of α-synuclein aggregation, we performed a genome-wide CRISPR interference screen in a human cell-based model. We discovered that inhibiting PIKfyve dramatically reduced α-synuclein aggregation induced with both recombinant α-synuclein fibrils and fibrils isolated from MSA patient brain. While PIKfyve inhibition did not affect fibril uptake or α-synuclein clearance or secretion, it reduced α-synuclein trafficking from the early endosome to the lysosome, thereby limiting fibril escape from the lysosome and reducing the amount of fibrils that reach cytosolic α-synuclein to induce aggregation. These findings point to the endolysosomal transport of fibrils as a critical step in the propagation of α-synuclein aggregation and a potential therapeutic target.

## INTRODUCTION

Neurodegenerative diseases are a pressing global public health challenge due to the lack of disease-modifying therapies. A major reason for the absence of effective treatments is likely our incomplete understanding of the molecular and cellular mechanisms underlying disease progression. A major advance in neurodegeneration research was the discovery that scrapie, an infectious neurodegenerative disease, is caused by the spread of protein aggregates via their prion forms.^1^ Prion proteins can template the conversion of healthy native proteins into an aggregated conformer that in turn continues to propagate. Since the discovery of prions, numerous independent studies suggest that the prion-like propagation of specific neuronal proteins such as tau and α-synuclein drive noninfectious neurodegenerative diseases such as Alzheimer’s disease or Parkinson’s disease.^2–4^ Our understanding of the mechanisms that control the prion-like spread of these proteins is incomplete. Fully elucidating these mechanisms systematically will further our understanding of disease progression and provide new therapeutic targets.

Here, we investigate the mechanisms underlying the prion-like spreading of α-synuclein aggregation with the goal to pinpoint potential therapeutic targets to inhibit this process. α-synuclein aggregation is a common hallmark of Parkinson’s disease, and is strongly correlated with neuronal death and subsequent cognitive and motor dysfunction.^5^ Copy number variations or point mutations in the α-synuclein gene have been linked to familial Parkinson’s Disease.^5^ α-synuclein aggregation occurs in other neurodegenerative diseases, broadly categorized as synucleinopathies, including multiple systems atrophy (MSA).^5^

α-synuclein aggregation and propagation can be modeled in cultured HEK293T cells that express fluorescently tagged versions of α-synuclein harboring patient-derived point mutations.^6–8^ In this system, addition of α-synuclein fibrils to the culture medium triggers the conversion of diffuse soluble α-synuclein into aggregated puncta. This cell-based model has enabled the characterization of α-synuclein species with seeding activity from post-mortem patient brains, further validating the physiological relevance of these cell-based models.^9–11^

These biosensors and others^12^ have been instrumental in detecting seeding activity of α-synuclein preparations. Importantly, cell-based models can also be used to discover new pathways that control propagation of α-synuclein aggregation. While previous studies sought to define the mechanisms of α-synuclein toxicity in yeast^13,14^ and mechanisms that regulate levels of α-synuclein in mammalian cells^15–17^, cellular pathways that control propagation of α-synuclein aggregation in mammalian cells have not been systematically characterized. We hypothesized that such pathways could encompass those that control uptake of α-synuclein fibrils, trafficking of fibrils through the endolysosomal system and escape to the cytosol, templated aggregation of soluble α-synuclein, and aggregate clearance (**Fig. 1A**).

**Fig. 1.**
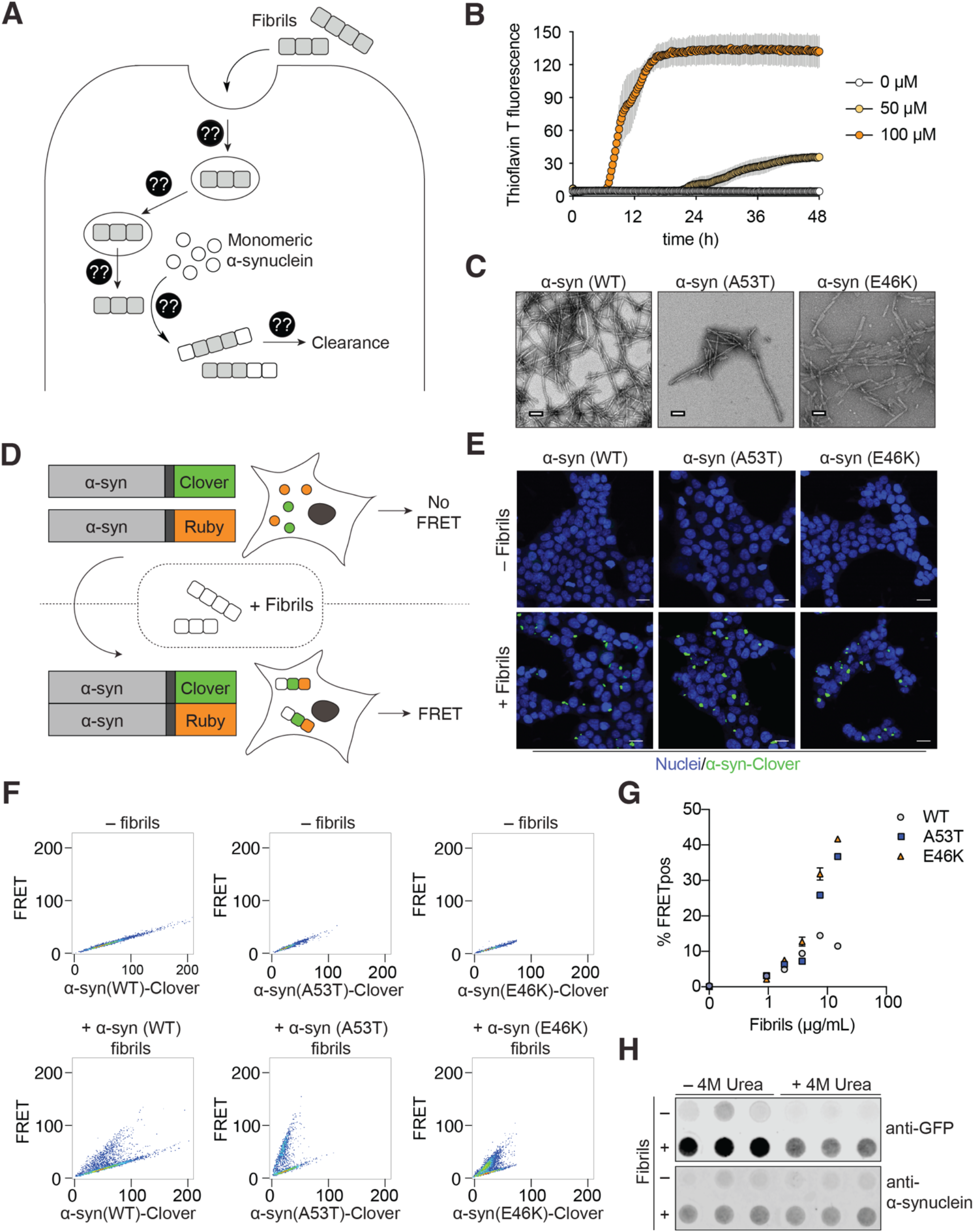
A cell-based model of prion-like spreading α-synuclein aggregation. (A) Diagram of cellular mechanisms that may control the prion-like spread of α-synuclein aggregation. Question marks represent unknown cellular mechanisms and potential therapeutic targets. (B) Vigorous agitation of recombinant α-synuclein monomers at 37°C induces fibrilization. Fibrilization is monitored by Thioflavin T fluorescence (excitation: 440 nm, emission: 485 nm). Error bars represent mean ± S.E.M from n = 4 technical replicates. (C) Representative negative stain electron micrographs α-synuclein fibrils formed from recombinant protein with the WT sequence or the Parkinson’s Disease-associated mutations A53T or E46K. Scale bar: 100 nm. (D) Schematic of the FRET-based reporter system used to monitor α-synuclein aggregation in HEK293T cells. In the absence of fibrils α-synuclein-Clover and α-synuclein-Ruby are monomeric. Exposure to α-synuclein fibrils induces aggregation of the reporter. During aggregation, the proximity of the fluorescently tagged α-synuclein can be detected by fluorescence microscopy or by flow cytometry. (E) Representative fluorescence microscopy images of reporter cells exposed to α-synuclein fibrils or lipofectamine alone. Blue: Draq5(nuclei). Green: α-synuclein-clover. Scale bar: 20 μm. (F) Representative flow cytometry plots of the WT, A53T and E46K FRET reporter cells after 24 h treatment with lipofectamine only (top row) or with α-synuclein fibrils (bottom row). (G) Quantification of % FRET-positive cells using flow cytometry across concentration ranges of α-synuclein fibrils. Error bars represent mean ± S.E.M from n = 3 technical replicates. (H) E46K FRET-reporter cells exposed to E46K fibrils were passed through a 0.45 μm nitrocellulose membrane after lysis and 10 min incubation in either TBS or 4M Urea. Insoluble material trapped on the membrane was quantified by immunoblotting against the clover tag to detect intracellular α-synuclein.

In order to discover cellular pathways that control spreading of α-synuclein aggregation, we applied our CRISPR interference-based (CRISPRi) genetic screening platform^18,19^ to a fluorescence resonance energy transfer (FRET) based model of α-synuclein aggregation in mammalian cells. Using this approach, we identified the phosphotidylinositol pathway as a critical regulator of the spreading of α-synuclein aggregation. Inhibition of the synthesis of multiple phosphatidylinositol species lowered aggregation. Specifically, inhibition of the kinase PIKfyve dramatically reduced α-synuclein trafficking through the endolysosomal system, thereby limiting the escape of α-synuclein seeds from lysosomal compartments and slowing subsequent templating of α-synuclein aggregation in the cytosol. These discoveries define mechanisms of α-synuclein trafficking that can be inhibited pharmacologically to reduce aggregation, highlighting their potential as therapeutic targets.

## RESULTS

### Cell-based model of prion-like propagation of α-synuclein aggregation

In order to identify cellular factors that control the prion-like propagation of α-synuclein aggregation (**Fig. 1A**), we established a cell-based model with a FRET readout. A similar FRET-based model had been previously developed by the Diamond Lab to monitor aggregation by microscopy and flow cytometry.^9^ Given the different aggregation propensities of wild-type α-synuclein and the Parkinson’s disease-associated A53T and E46K point mutations, we generated three versions of this cell-based system: wild-type, A53T and E46K.

First, we generated recombinant α-synuclein fibrils to serve as aggregation seeds. We purified monomeric human α-synuclein from *Escherichia coli* and induced fibrillization by vigorous agitation at 37 ºC. We verified the generation of amyloid fibrils by the increase in Thioflavin T fluorescence (**Fig. 1B**) and negative stain EM (**Fig. 1C**). These fibrils were subsequently sonicated into smaller seeds.

Next, we co-expressed two versions of full-length human α-synuclein fused to the fluorescent proteins mClover2 and mRuby2, respectively, in HEK293T cells. We reasoned that upon triggering aggregation with α-synuclein fibrils added to the cells, the monomeric fusion proteins would come within close proximity to each other and generate a detectable FRET signal (**Fig. 1D)**.

Fluorescent constructs were stably introduced to HEK293T cells by lentivirus. We then selected monoclonal lines based on high expression of the fusion proteins and dynamic range of the FRET signal. In the absence of fibrils, these reporter cells showed diffuse intracellular fluorescence without visible aggregates when monitored by fluorescence microscopy (**Fig. 1E**) and appeared as a single population when FRET levels were monitored by flow cytometry (**Fig. 1F**). Upon exposure to recombinant α-synuclein fibrils complexed with lipofectamine, we observed that the fluorescent reporter constructs formed visible puncta by fluorescent microscopy (**Fig. 1E**), and a second distinct FRET-positive population of cells emerged by flow cytometry (**Fig. 1F**), indicating aggregation of the intracellular α-synuclein constructs. We found that all three fibril types triggered aggregation in a dose-dependent manner when added to cell types expressing the reporters with the corresponding sequences (WT, A53T or E46K), as quantified by the percentage of FRET-positive cells (**Fig. 1G**). The E46K α-synuclein FRET reporter line reproducibly generated the highest percentage of FRET-positive cells, and generated detergent and urea resistant aggregates (**Fig. 1H**), making it an ideal cell line to use in a genome-wide genetic screen to identify cellular factors controlling α-synuclein aggregation.

### Genome-wide screen for modifiers of prion-like propagation of α-synuclein aggregation

To enable a genetic screen for modifiers of propagation of α-synuclein aggregation, the cell lines described above also expressed catalytically inactive Cas9-BFP-KRAB (dCas9-BFP-KRAB) fusion protein. dCas9-BFP-KRAB can be directed by single guide RNAs (sgRNAs) to knock down a specific gene of interest,^20^ enabling large-scale genetic screens in mammalian cells.^18^

The screening strategy is illustrated in **Figure 2A**. We transduced these cells with pooled lentiviral libraries containing sgRNAs targeting every protein-coding gene in the genome and nontargeting control sgRNAs. Cells transduced with sgRNAs were exposed to recombinant α-synuclein fibrils at concentrations that yielded the half-maximal percentage of FRET-positive cells, thereby maximizing the dynamic range for detecting cellular factors that either increase or decrease α-synuclein aggregation. FRET-negative and FRET-positive cell populations were separated by FACS. Sufficient numbers of cells were collected from each population to achieve at least 500× average representation of cells per sgRNA elements in the library. Genomic DNA was isolated, sgRNAs were PCR-amplified, and frequencies for each sgRNA in each population were quantified by next-generation sequencing. We evaluated the effect of gene knockdown on the formation of α-synuclein aggregates using our previously described bioinformatics pipeline.^18,21–23^

**Fig. 2.**
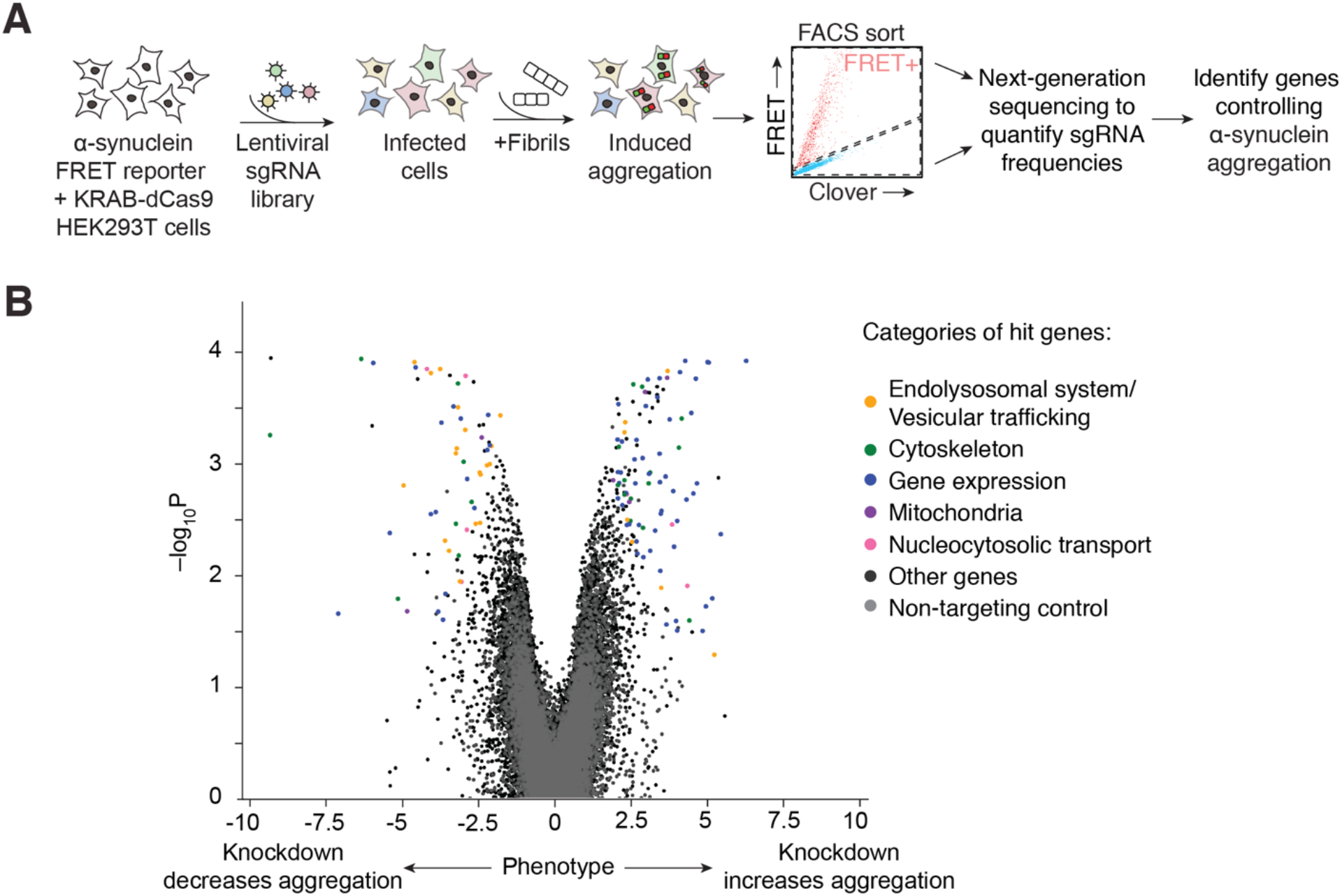
A genome-wide CRISPRi screen for cellular factors controlling α-synuclein aggregation. (A) Schematic for a pooled FRET-based CRISPRi screen. FRET reporter cells stably expressing CRISPRi machinery (dCas9-BFP-KRAB) were transduced with pooled lentiviral libraries encoding sgRNAs targeting every protein-coding gene in the genome. Infected cells were then selected, expanded and exposed to α-synuclein fibrils. 24 hours later, cells were sorted into FRET-negative and FRET-positive populations by FACS. sgRNA-encoding cassettes were amplified from the genomic DNA of each population and sgRNA frequencies were quantified by next generation sequencing in order to identify genes that control α-synuclein aggregation. (B) Volcano plot of genes showing categories of major hit genes from the primary genome-wide screen.

The genome-wide screen uncovered genes knockdown of which either increased or decreased α-synuclein aggregation (**Table S1**). Many of the top hits could be assigned to one of the following biological categories: Endolysosomal system / vesicular trafficking, cytoskeleton, gene expression, mitochondria, nucleocytosolic transport (**Fig. 2B, Table S1**).

Next, we selected hits from the primary genome-wide screen, which was performed with the α-synuclein (E46K) FRET-reporter, for further characterization in secondary screens. These hits were first selected from genes with an FDR < 0.1 and further narrowed to exclude heavily redundant genes specific to a biological pathway, resulting in a focused library targeting 249 genes (**Table S2**). Using this focused library, we conducted secondary screens with all three versions of the α-synuclein reporter. The hits targeted by this focused retest library revalidated in all three cell lines robustly with minimal differences between the α-synuclein variants (**Fig. 3A, Fig. S1, Table S2**).

**Fig. 3.**
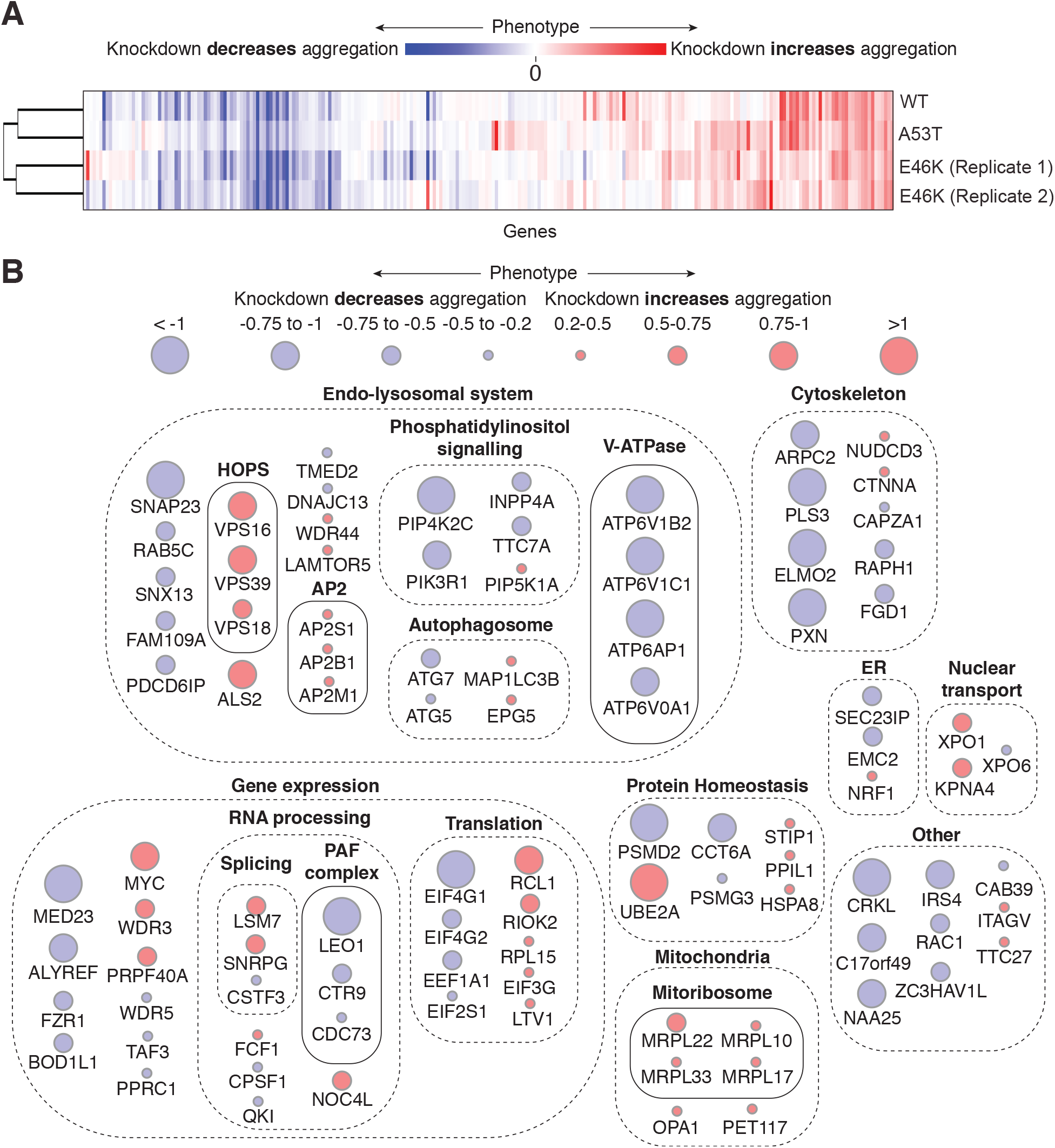
Focused revalidation CRISPRi screen for cellular factors controlling α-synuclein aggregation. (A) Heat map summarizing the relative phenotypes from four focused secondary screens: WT, A53T and E46K reporters Genes selected for these focused screens were top representative hits from the primary genome-wide CRISPRi screen. (B) Overview of top hits from these focused screens, and the biological pathways in which they act.

Genes that reproduced strongly in the WT, A53T, and E46K cell lines are displayed in **Figure 3B**, grouped by their biological pathways. The knockdown of many genes in the endolysosomal system dramatically reduced aggregation. One major pathway stood out for its strong impact on α-synuclein aggregation: the phosphatidylinositol pathway, a major signaling pathway that regulates, among other processes, endolysosomal trafficking and autophagy. We decided to prioritize this pathway for further characterization because knockdown of several factors in this pathway robustly reduced aggregation, making it a potential therapeutic target.

### Pharmacological inhibition of PIKfyve and other phosphatidylinositol kinases reduces α-synuclein aggregation

In order to comprehensively test which members of the phosphatidylinositol pathway control aggregation, we selected a panel of small molecule inhibitors of major kinases involved in this pathway (**Fig. 4A, Fig. S2**). Interestingly, the majority of these inhibitors blocked aggregation robustly (**Fig. 4A**). In particular, apilimod, an inhibitor of PI-3P-5-kinase (PIKfyve), which phosphorylates the D-5 position in phosphatidylinositol-3-phosphate (PI3P) to yield phosphatidylinositol 3,5-bisphosphate (PI(3,5)P2),^24^ dramatically reduced aggregation with IC50 values below 25 nM (**Fig. 4B).** To confirm these results genetically, we cloned two individual sgRNAs each targeting PIKfyve and validated that PIKfyve knockdown in our FRET reporter lines also reduced aggregation (**Fig. 4C**, **Table S2**) but to a lesser extent compared to the pharmacological inhibitors. PIKfyve inhibition also reduced the amount of insoluble α-synuclein, measured by a filter trap assay, in a dose-dependent manner (**Fig. 4D,E**).

**Fig. 4.**
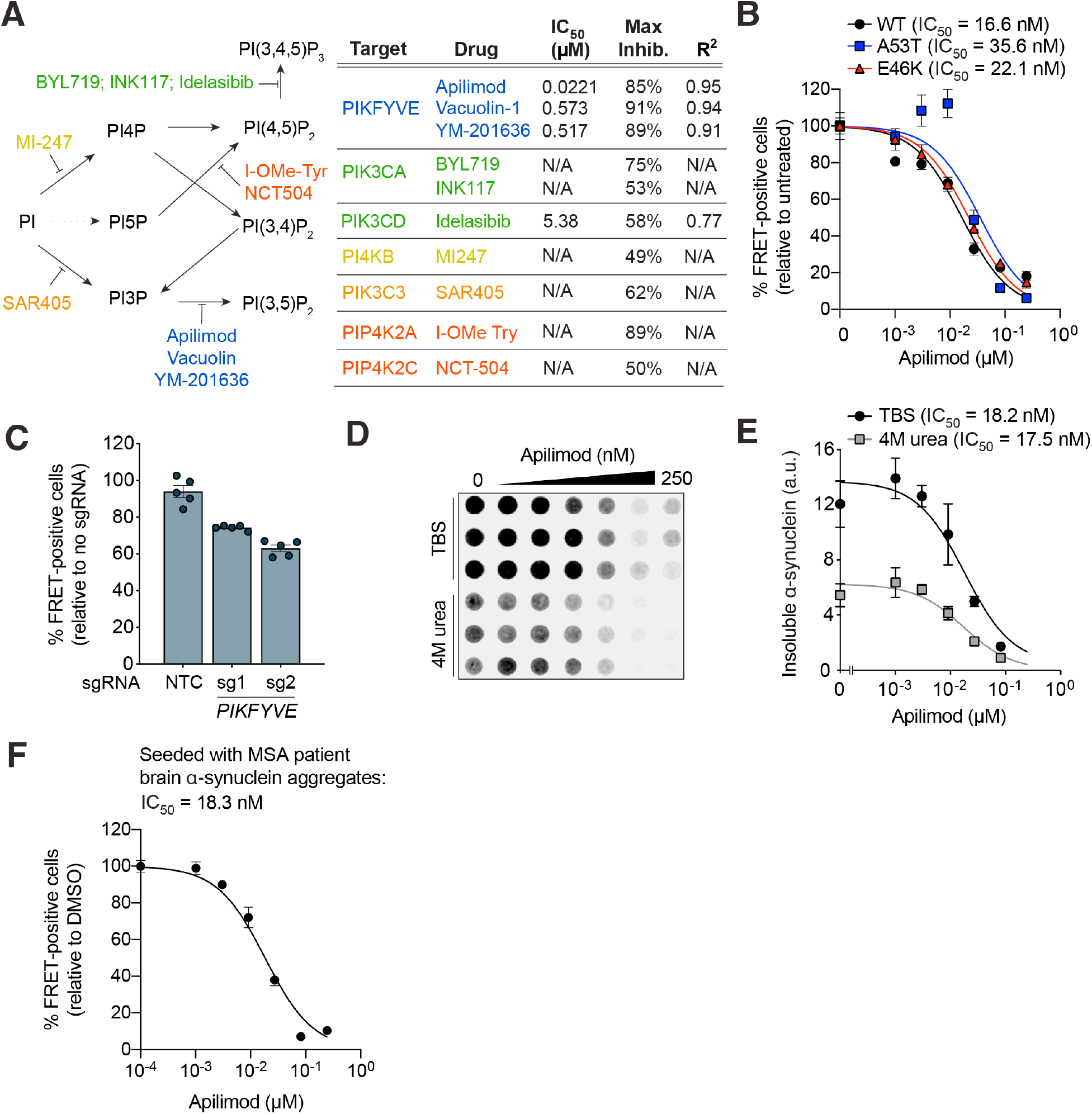
Inhibition of PIKFYVE decreases α-synuclein aggregation in FRET reporter cells seeded with fibrils. (A) Pharmacological inhibition of several kinases in the phosphatidylinositol pathway decreases α-synuclein aggregation. FRET reporter cells were co-treated with fibrils and various concentrations of each kinase inhibitor and collected for flow cytometry after 24 hours (also in Supplemental Figure 1). (B) Dose-dependent decrease of aggregation in FRET reporter cells treated with the PIKFYVE inhibitor apilimod. Error bars represent mean ± s.e.m from n = 3 technical replicates. (C) FRET reporter cells were transduced with individual sgRNAs targeting PIKFYVE or a non-targeting control sgRNA (NTC) and aggregation was measured 5 days after transduction. Error bars represent mean ± S.E.M from n = 3 technical replicates. (D) Apilimod decreases insoluble intracellular α-synuclein. FRET-reporter cells exposed to increasing apilimod concentrations were passed through a 0.45 μm nitrocellulose membrane after lysis and 10 min incubation in either TBS or 4M Urea. Insoluble material trapped on the membrane was quantified by immunoblotting against the clover tag to detect intracellular α-synuclein. (E) Quantification of Fig. 3D. Immunoblot signal was normalized to lysate protein concentration as determined by a BCA assay. Error bars represent mean ± S.E.M from n = 3 technical replicates. (F) Dose-dependent decrease of aggregation trigged with MSA patient brain-derived fibrils derived with the PIKFYVE inhibitor apilimod. Error bars represent mean ± s.e.m from n = 3 technical replicates.

Recently, α-synuclein fibrils isolated from post-mortem brains from patients with a synucleinopathy called multiple system atrophy (MSA) were shown to trigger aggregation in cellular and mouse models of α-synuclein propagation.^6,9,10^ We tested the effect of PIKFyve inhibition on α-synuclein aggregation triggered by α-synuclein fibrils isolated from deceased MSA patient brain samples. We found that PIKYVE inhibition decreased α-synuclein aggregation in our FRET reporter line seeded with patient brain-derived α-synuclein by flow cytometry (**Fig. 4F**).

### PIKfyve inhibition does not increase autophagic flux, α-synuclein secretion, nor decrease fibril uptake

We next investigated the mechanism by which PIKfyve inhibition reduces α-synuclein aggregation. Given the importance of autophagy for the clearance of aggregated proteins, including α-synuclein,^25^ we hypothesized that PIKfyve might reduce aggregation by increasing autophagic flux. We first evaluated the effect of PIKfyve inhibition on autophagy using a LC3 cell-based reporter. This reporter consists of a fusion construct of GFP-LC3 and RFP-LC3ΔG that is cleaved by ATG4 during autophagy. GFP-LC3 is degraded by autophagy whereas RFP-LC3ΔG remains in the cytosol as an internal control. Autophagic flux can then be quantified by flow cytometry as the ratio of RFP to GFP.^26^ Surprisingly, we discovered that PIKfyve inhibition decreased, rather than increased autophagic flux (**Fig. 5A**) in the presence of torin, a pharmacological activator of autophagy.

**Fig. 5.**
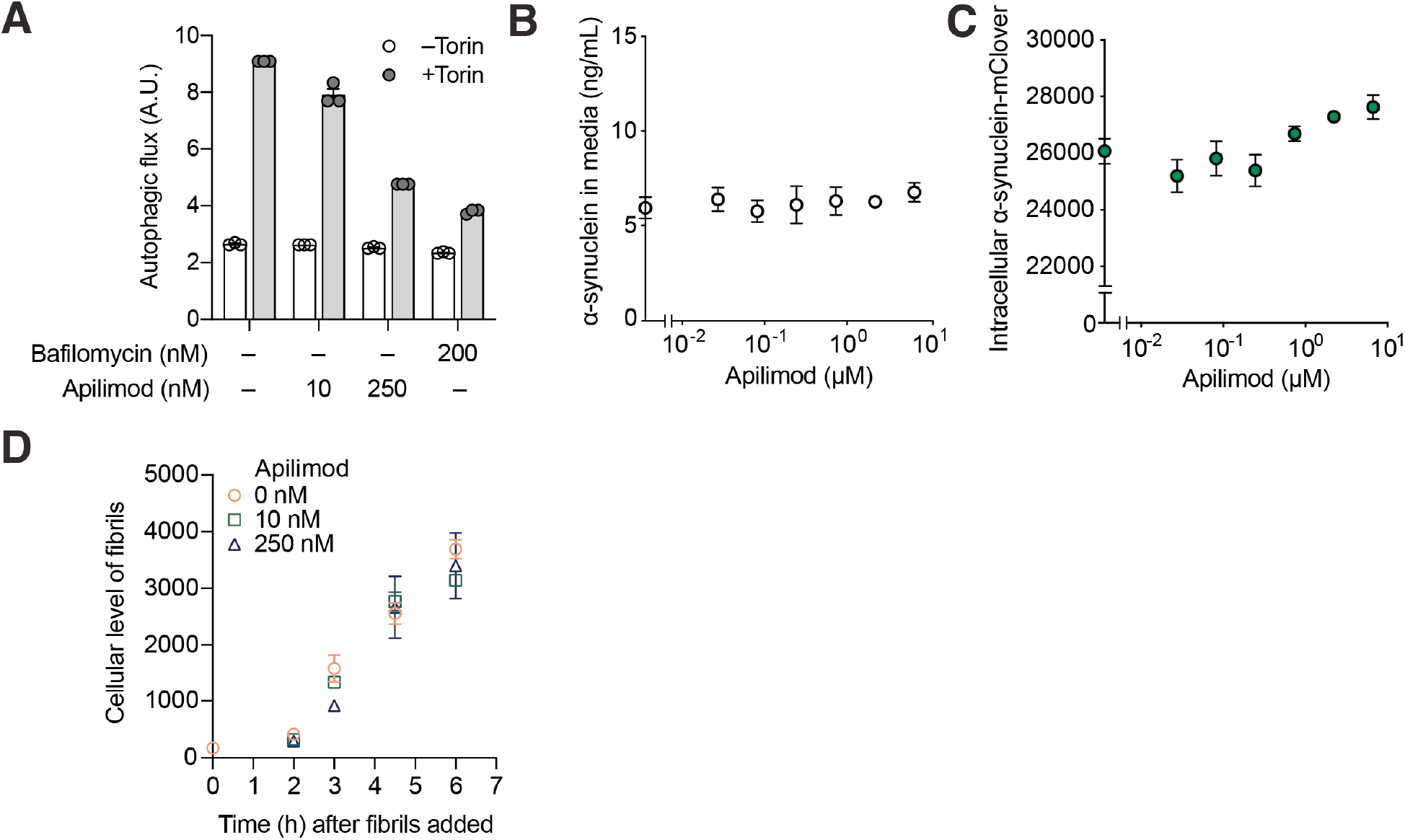
PIKFYVE inhibition decreases autophagic flux but does not affect α-synuclein export or fibril uptake. (A) CRISPRi-HEK293T cells expressing mCherry-GFP-LC3 were treated with either bafilomycin or apilimod in the presence and absence of torin. Changes in the ratio of GFP/mCherry was monitored by flow cytometry. Error bars represent mean ± S.E.M from n = 3 technical replicates. (B) α-synuclein and mClover protein detected in the media by ELISA. Error bars represent mean ± S.E.M from n = 3 technical replicates. (C) PIKFYVE inhibition does not impact steady-state levels of the α-synuclein-Clover2 construct in FRET reporter cells, as quantified by flow cytometry. Error bars represent mean ± S.E.M from n = 3 technical replicates. (D) PIKFYVE inhibition does not impact uptake of Alexa-555 tagged fibrils. Error bars represent mean ± S.E.M from n = 3 technical replicates.

Previous studies have suggested that the inhibition of autophagy can promote the exocytosis of α-synuclein as an alternative clearance mechanism.^27^ We therefore hypothesized that PIKfyve inhibition may lower levels of aggregated α-synuclein by promoting export of intracellular α-synuclein. To test this hypothesis, we performed an ELISA to detect the level of α-synuclein-Clover in the cell culture media after triggering aggregation (**Fig. 5B**). PIKfyve inhibition did not cause an increase in the amount of α-synuclein-Clover in the media, and there was no decrease in intracellular α-synuclein-Clover levels (**Fig. 5C**), suggesting that an export mechanism could not explain the dramatic decrease in aggregation during PIKfyve inhibition.

We then evaluated if PIKFyve inhibition reduced aggregation by blocking the initial fibril uptake into cells. By measuring the uptake of fluorescently tagged fibrils at different apilimod concentrations, we determined that PIKFyve inhibition did not alter fibril uptake (**Fig. 5D**).

### PIKfyve inhibition interferes with α-synuclein fibril trafficking to the lysosome and lysosomal rupture

Given the role of PIKfyve at multiple stages of the endolysosomal pathway,^28^ we sought to investigate if PIKfyve inhibition could alter fibril trafficking. We monitored fibril transport in relationship to the early endosome marker EEA1 (**Fig. 6A**) and the late endosome/lysosome marker LAMP1 (**Fig. 6B**) at 30 min, 1 h, 2h, 4h, and 24 h after fibril exposure. In the absence of PIKfyve inhibition, fibrils are not detected in EEA1-positive compartments at these time points (**Fig. 6A**), but accumulate in LAMP1-positive compartments as early as 30 min after fibril exposure (**Fig. 6B**). In the presence of apilimod, which resulted in enlargement of both EEA1 and LAMP1 vesicles, fibrils colocalized with EEA1 vesicles at the 1-hour timepoint, dramatically reducing the number of fibrils that reach LAMP1 vesicles at early and late timepoints. These results suggest that PIKfyve inhibition reduces fibril trafficking from the early to late endosome/lysosome.

**Fig. 6.**
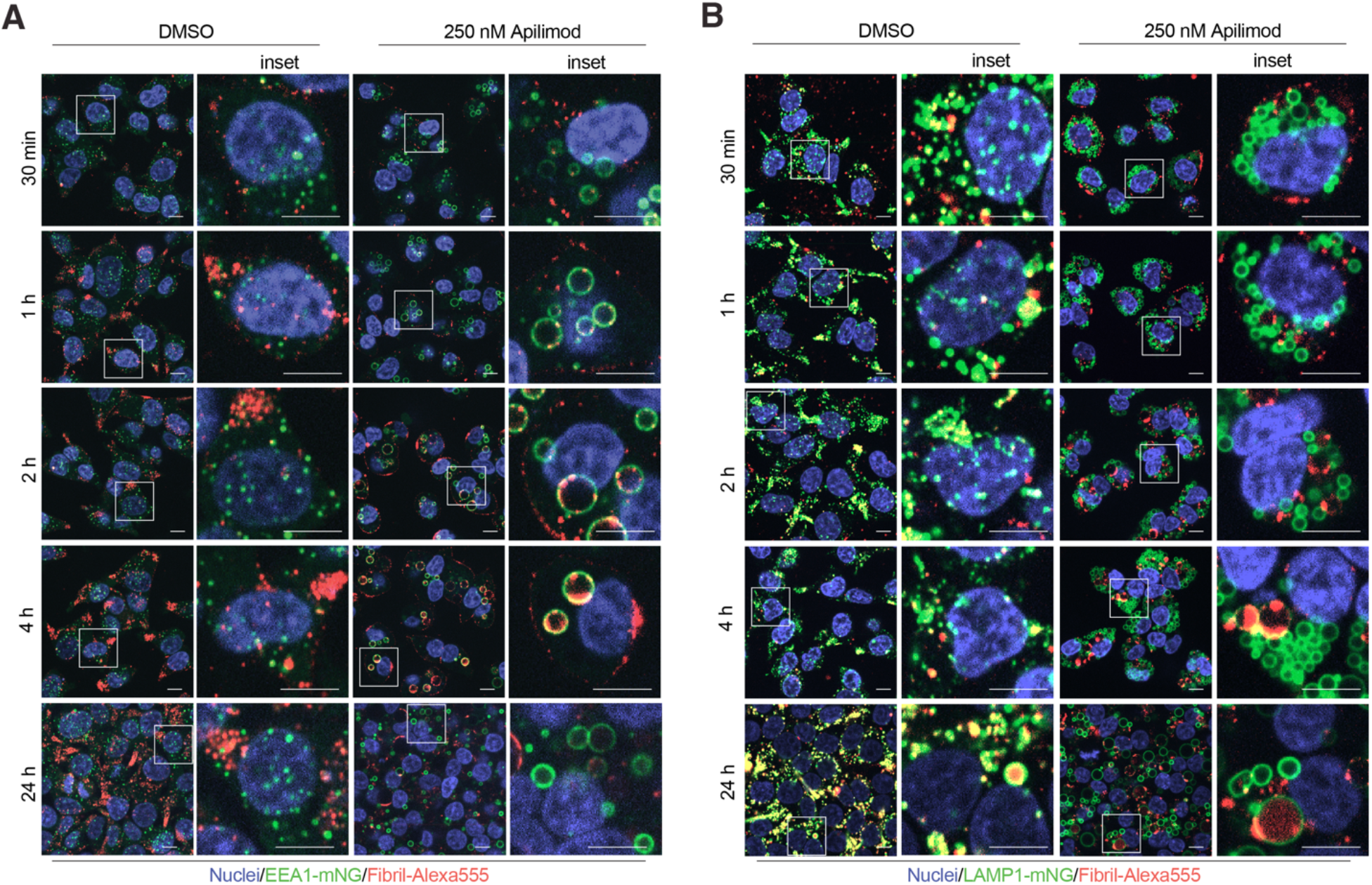
PIKFYVE inhibition interferes with α-synuclein fibril trafficking to the lysosome. Representative fluorescence microscopy images of cells in which the early endosomal marker EEA1 (A) or the lysosomal marker LAMP1 (B) were endogenously labeled with the split-mNeonGreen system and exposed to fibrils tagged with Alexa555 in the presence and absence of Apilimod. Timepoints were acquired at 30 min, 1 h, 2h, 4h, and 24 h after fibril addition. Scale bar: 5 μm.

α-synuclein fibrils have been previously shown to cause endolysosomal damage,^29^ leading us to hypothesize that PIKFyve inhibition may reduce α-synuclein aggregation by reducing fibril-induced rupture of the lysosome and subsequent escape of fibrils to the cytosol. We tested this hypothesis by monitoring the formation lysosomal damage using a cytosolic GFP fusion of galectin 3 (GAL3), a lectin that binds β-galactosides and forms puncta when these sugars are exposed on damaged endolysosomes.^30^ Consistent with previously reported studies^29^ α-synuclein fibrils caused endolysomal damage. Pharmacological and genetic PIKFyve inhibition dramatically reduced α-synuclein fibril-induced endolysomal damage (**Fig. 7A-C**).

**Fig. 7.**
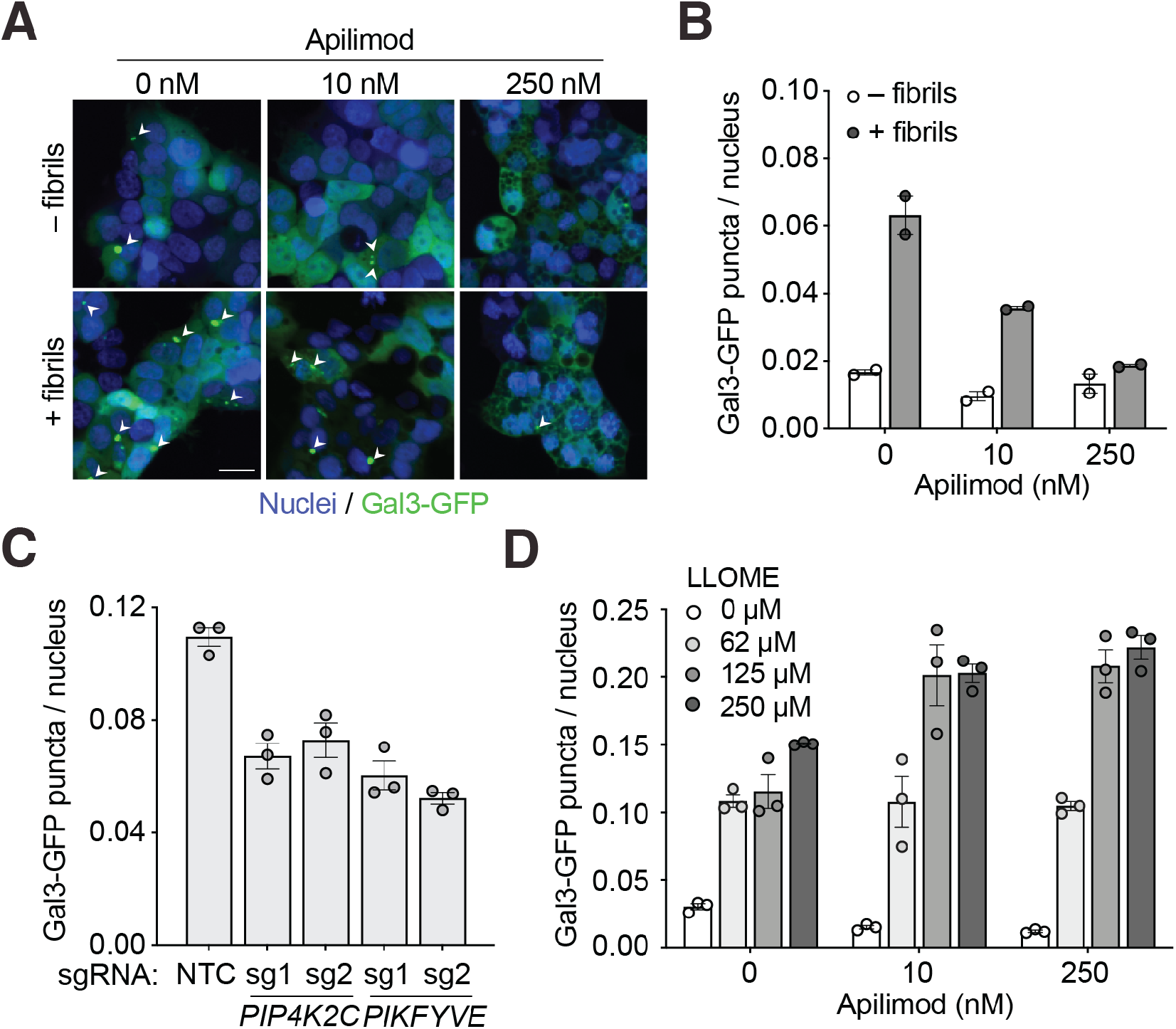
PIKFYVE inhibition reduces fibril-induced lysosomal rupture. (A) Representative fluorescence microscopy images of CRISPRi-HEK293T cells expressing an EGFP-Galectin3 (EGFP-Gal3) reporter treated with apilimod. EGFP-Gal3 reporter cells were co-treated with fibrils and apilimod or DMSO 24 hours prior to imaging. Nuclei were counter-stained with Draq5. Scale bar: 20 μm. (B) Quantification of EGFP-Gal3 puncta divided by number of nuclei in fluorescence microscopy images shown in (A). Error bars represent mean ± S.E.M for *n* = 2 technical replicates (with at least 300 nuclei per image). (C) EGFP-Gal3 reporter cells were transduced with individual sgRNAs targeting PIP4K2C, PIKFYVE or a non-targeting control sgRNA (NTC). 5 days after transduction fibrils were added to EGFP-Gal3 reporter cells and puncta formation was observed by microscopy. Error bars represent mean ± S.E.M for *n* = 3 technical replicates (with at least 300 nuclei per image). (D) EGFP-Gal3 reporter cells were treated with various concentrations of LLOME and apilimod and 24 h after treatment puncta formation was observed by microscopy. Error bars represent mean ± S.E.M for *n* = 3 technical replicates (with at least 300 nuclei per image).

Given the strong reduction of lysosomal damage upon PIKFYVE inhibition, we asked whether PIKFyve inhibition generally protects lysosomes from damage, or whether the effect is specific to damage induced by α-synuclein fibrils. To address this question, we treated cells with leucyl-leucyl-*O*-methyl ester (LLOME), a compound that accumulates in the lumen of acidified organelles, rapidly forms membranolytic polymers after cleavage by cathepsin C,^30,31^ thereby causing direct lysosomal rupture. We confirmed that LLOME damages endolysosomal membranes in our cell line using the GAL3-GFP reporter (**Fig. 7D**). Apilimod did not protect cells from LLOME-induced lysosomal damage, but rather sensitized them (**Fig. 7D**), indicating that its protective effect in the presence of α-synuclein fibrils is due to inhibition of fibrils reaching the lysosome, as opposed to a direct effect on lysosomal integrity.

These results together suggest that PIKfyve controls fibril trafficking from the early to late endosome, and that the inhibition of PIKfyve decreases the amount of fibrils reaching lysosomes. The resulting lysosomal rupture limits fibrils from templating aggregation of soluble α-synuclein in the cytosol, inhibiting α-synuclein aggregate spreading (**Fig. 8**).

**Fig. 8.**
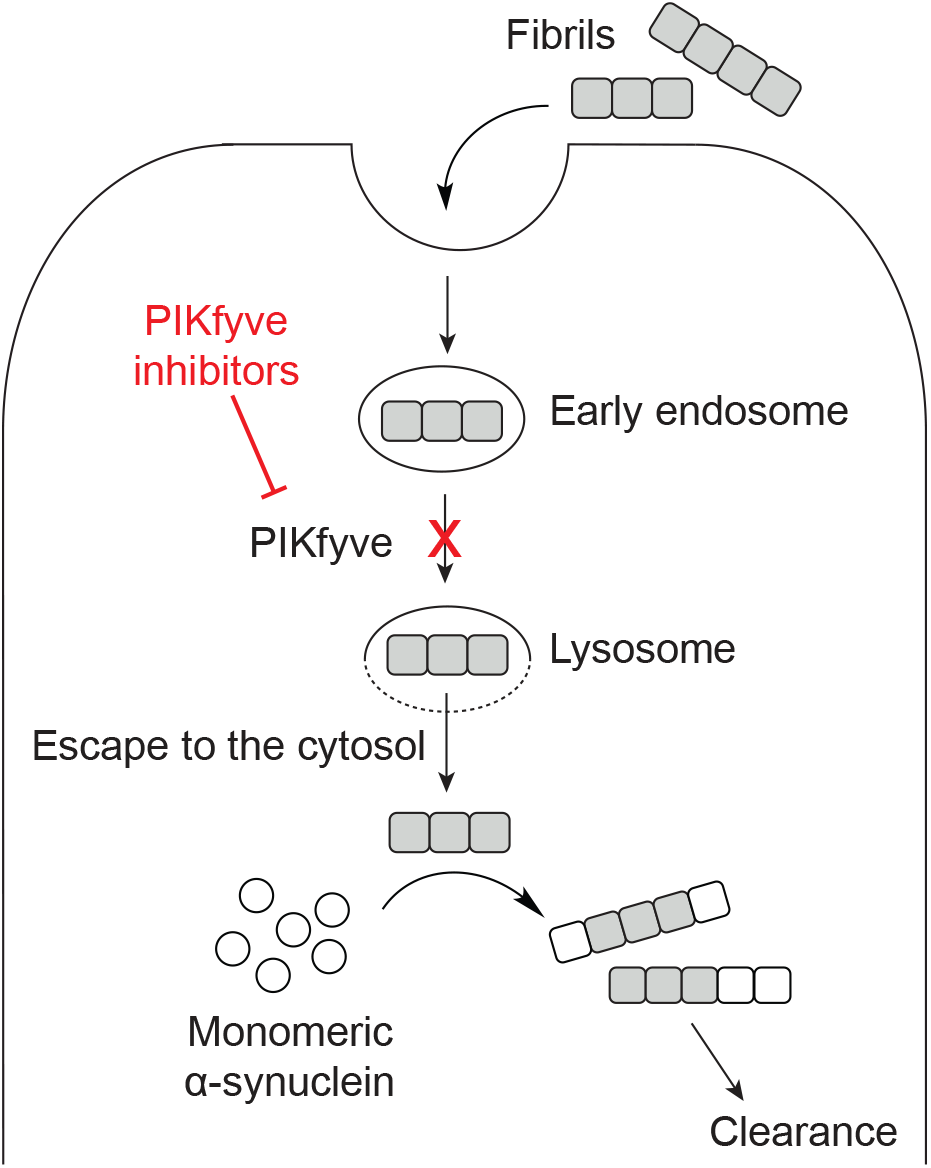
Model. PIKfyve inhibition reduces propagation of α-synuclein aggregation in cells by limiting fibril access to the lysosome and lysosomal escape.

## DISCUSSION

We used our CRISPRi-based genetic screening platform to discover new mechanisms controlling prion-like spreading of α-synuclein aggregation in a cell-based model, and potential therapeutic targets to block this process. Importantly, we discovered that the inhibition of PIKfyve decrease aggregation by blocking fibril trafficking to lysosomes, thereby decreasing the endolysosomal escape of α-synuclein fibrils.

Since our observations were made in an *in-vitro* model of α-synuclein aggregation, it remains to be tested whether inhibition of PIKfyve may reduce spread and propagation of α-synuclein in the context of neurodegenerative diseases *in vivo*. Beyond its potential to reduce prion-like spreading of protein aggregation, inhibition of PIKfyve has been shown to have other beneficial effects in neurodegeneration as well. The inhibition of PIKfye has been shown to be beneficial in mouse models of amyotrophic lateral sclerosis and frontotemporal dementia induced by repeat expansions in C9orf72 by reversing endolysosomal defects.^32^

Intriguingly, PIKfyve inhibition has been shown to block infection with SARS-CoV-2 *in vitro*, likely by a mechanism similar to the one we have delineated here for α-synuclein fibrils^33^. Apilimod is currently evaluated in a clinical trial for efficacy in COVID-19 patients (NCT04446377). This suggests that PIKfyve inhibition can block endolysosomal trafficking of both pathogenic protein aggregates and viruses, and effective inhibitors could have a broad spectrum of clinical uses.

Our genome-wide screen also pointed to additional pathways relevant for prion-like propagation of α-synuclein aggregation, including other trafficking factors, such as the Parkinson’s risk gene DNAJC13. Another major class of hits is related to gene expression (including translation and ribosome biogenesis). These hits, however, may be pleiotropic as they likely affect expression of many genes.

We had expected to discover a larger network of molecular chaperones that regulate α-synuclein aggregation. Numerous *in vitro* and *in vivo* studies point to chaperones and their co-chaperones that strongly control aggregation and may be important therapeutic targets for synucleinopathies.^34,35^ The knockdown of individual chaperones did not have a major impact on α-synuclein aggregation in our genetic screen. This could be due to the redundancy of molecular chaperones in cells and would be better addressed through gain-of-function studies.

Compared to previous genetic screens focused primarily on toxicity, our work identifies cellular mechanisms upstream of protein aggregation. Despite our ability to identity components of the endolysosomal pathway responsible for trafficking, the use of Lipofectamine may prevent our detection of some very early cellular components important for α-synuclein uptake, such as the receptor Lag3^36^ and other early endocytic factors. Future screens in iPSC-derived neurons using our newly developed platform^23^ may uncover additional mechanisms critical to both α-synuclein aggregation and toxicity in specific neuronal subtypes.

## MATERIALS AND METHODS

### Purification, fibrillization, characterization and labelling of recombinant α-synuclein

Human WT, A53T and E46K α-synuclein proteins were expressed in BL2-Gold(DE3) competent cells (Agilent, Cat #230132). Protein expression was induced at log phase with 200 μM IPTG for 16 hours at room temperature. Cells were lysed in TE buffer (1 mM EDTA in 10 mM Tris pH 8.0 containing protease inhibitors) using sonication and the suspension was boiled for 30 min. The lysate was clarified and α-synuclein was subsequently precipitated with ammonium sulfate (361 mg/mL). The resulting pellet was resuspended in 50 ml of 25 mM sodium phosphate buffer and run over a HiTrapQ-HP 5 mL column (Cytiva, Cat# 17115401). α-synuclein was eluted with elution buffer (25 mM sodium phosphate buffer with 1 M NaCl over a gradient). Fractions containing α-synuclein confirmed by Coomassie-stained SDS-PAGE were dialyzed into PBS and concentrated with an Amicon Ultra-15 centrifugal 3 kDa MWCO filter (Millipore, Cat# UFC900324), filter sterilized with using a Millex-GV syringe filter unit (Millipore, Cat# SLGV033RB), and snap frozen in PBS at −80°C.

Aggregation was induced by agitating 5 mg/mL α-synuclein at 37 °C for 7 days 1200 rpm (VWR, Cat#12620-942). Fibril formation was evaluated by electron microscopy (see next section for details). Resulting fibrils were diluted in PBS to 0.25 mg/mL transferred to sonication tubes (Diagenode Cat# C30010011) and sonicated (Diagenode Bioruptor Pico, Cat# B01060010) to generate fibril seeds for cell-based assays.

In order to monitor the kinetics of α-synuclein fibril formation *in vitro* 20 μM of Thioflavin T was incubated with 5 mg/mL WT, A53T, or E46K monomer for 3 days at 37 °C with continuous shaking and monitored via Thioflavin T fluorescence (excitation=444 nm, emission=485 nm, cutoff=480 nm) in a Spectramax M5 microplate reader (Molecular Devices). Readings were taken every 15 min.

To generate fluorescently labeled fibrils, 0.6 μl of 10 mg/ml of Alexa Fluor 555 (ThermoFisher, Cat# A37571), 100 μl of 0.25 mg/ml of fibrils, and 9.4 μl of 1 M sodium bicarbonate were mixed at room temperature in the dark for 1 hour. Labeled α-synuclein fibrils were separated from unlabeled dye with using a Zebra 7k MWCO spin desalting column (ThermoFisher, Cat# 89882).

### Preparation of fibrils from multiple system atrophy patient brains

Phosphotungstic acid (PTA) was performed as described (^6,37,38^). Briefly, 10% (w/v) brain homogenate was incubated in 2% (v/v) sarkosyl (Sigma, 61747-100ML) and 0.5% (v/v) benzonase (Sigma, # E104-25KU) at 37 °C with constant agitation (1,200 rpm) in an orbital shaker for 2 hours. 10% Sodium PTA (Sigma, P6395-100G) was dissolved in water, and the pH was adjusted to exactly 7.0. The 10% NaPTA solution was added to the solution to a final concentration of 2% (v/v), which was agitated (1,200 rpm) in an orbital shaker overnight. The sample was centrifuged at 16,000 × *g* for 30 min at room temperature, and the supernatant was carefully removed. The pellet was resuspended in 2% (v/v) sarkosyl in DPBS and 2% (v/v) PTA in water, pH 7.0. The sample was again incubated for at least 1 hour before a second centrifugation. The supernatant was removed, and the pellet was resuspended in DPBS using 10% of the initial starting volume. This suspension was diluted 1:20 in DPBS before use in cell-based assays.

### Electron microscopy

α-synuclein fibrils were negatively stained with 0.75% uranyl formate (pH 5.5–6.0) on thin amorphous carbon-layered 400-mesh copper grids (Pelco Cat# 1GC400). 5 μl of 5 mg/mL fibrils was applied to the grid for 1 min before removing the droplet with Whatman paper, followed by two washes with 5 μl of ddH2O and two applications of 5 μl of uranyl formate removed by vacuum. Grids were imaged at room temperature using a Fei Tecnai 12 microscope operating at 120 kV. Images of WT and A53T fibrils were captured on a US 4000 CCD camera at x44117 resulting in a sampling of 3.4 Å/pixel. E46K fibrils were imaged on the microscope at x57692 resulting in a sampling of 2.6 Å/pixel.

### Plasmid and library design

Plasmids for the FRET-aggregation reporters were generated by subcloning full length human α-synuclein PCR-amplified from pCAGGS-aSyn-CFP (a gift from Robert Edwards and Ken Nakamura)^39^ to the C-terminal Clover2 or mRuby2 into the lentiviral expression vector pMK1200 (Addgene #84219)^21^ under the control of the constitutive EF1A promoter, to obtain pSKS40 and pSKS41, respectively. Point mutations were generated using QuikChange II site-directed mutagenesis kits (Agilent, Cat# 200521) according to the manufacturers protocols in pSKS40 and pSKS41 to generate α-synuclein(A53T)-mClover2 (pSKS42), α-synuclein(E46K)-mClover2 (pSKS44), α-synuclein(A53T)-mRuby2 (pSKS43), synuclein(A53T)-mRuby2 (pSKS45).

The α-synuclein(WT) bacterial expression plasmid was a gift from Stanley Prusiner. Point mutations were generated using QuikChange II site-directed mutagenesis kit to generate α-synuclein(A53T) (pSKS28) and α-synuclein(E46K) (pSKS39) plasmids for bacterial expression. Pooled CRISPRi sgRNA libraries genome-wide libraries were previously designed and validated.^40^

For generation of individual sgRNAs for focused retest screens, pairs of oligonucleotides (IDT) were annealed and ligated into our lentiviral sgRNA expression vector.

For stable expression of the CRISPRi machinery, we established lentiviral dCas9 expression vectors.^41^

The autophagy reporter plasmid pRT48 (CAG-GFP-LC3-RFP-LC3∆G) was generated by subcloning the GFP-LC3-RFP-LC3∆G-IRES-PuroR cassette from pMRX-IP-GFP-LC3-RFP-LC3∆G (a gift from Noboru Mizushima and Addgene plasmid # 84572) to a lentiviral backbone downstream of a CAG promoter.

### Cell culture and cell line generation

HEK293T cells were cultured in DMEM supplemented with 10% fetal bovine serum (Seradigm Cat# 97068-085, lot 076B16), Pen/Strep (Life Technologies, Cat# 15140122), and L-glutamine (Life Technologies, Cat# 25030081). All cells were grown in a tissue culture incubator (37 °C, 5% CO2) and regularly tested for mycoplasma contamination.

To introduce CRISPRi functionality to HEK293T cells, pHR-SFFV-dCas9-BFP-KRAB (Addgene #46911),^20^ a gift from Stanley Qi and Jonathan Weissman was stably expressed in HEK293T cells, and a monoclonal cell line (cSKS197) was selected and validated as previously described.^23^

To generate the three FRET reporter lines, cSKS197 cells were infected with lentivirus from plasmids pSKS40/pSKS41, pSKS42/pSKS43, pSKS44/pSKS45, and cells with the highest dynamic FRET signal 24 hours after seeding with fibrils were selected.

LAMP1 and EEA1 endolysosomal markers were endogenously labeled using the split-mNeonGreen2 system.^42^ Briefly, synthetic guide RNAs (IDT) were first complexed *in vitro* with purified *Streptococcus pyogenes* Cas9 protein (UC Berkeley Macrolab). Cas9/RNA complexes were then mixed with ssDNA oligo donors (IDT) and nucleofected (Lonza Cat# AAF-1002B, Amaxa program CM-130) into HEK293T cells stably expressing SFFV-mNeonGreen21–10. Fluorescent cells were selected by flow cytometry. Sequences for CRISPR RNA and donors used are listed as follows: LAMP1 (C-term mNG11): crRNA sequence, 5′-GTGCACCAGGCTAGATAGTC-3′; donor oligonucleotide sequence, 5′-CCCAGAGAAAGGAACAGAGGCCCCTGCAGCTGCTGTGCCTGCGTGCACCAGGCTACATCATATCGGTAAAGGCCTTTTGCCACTCCTTGAAGTTGAGCTCGGTACCACTTCCTGGACCTTGAAACAAAACTTCCAATCCGCCACCGATAGTCTGGTAGCCTGCGTGACTCCTCTTCCTGCCGACGAGGTAGGCGATGAGG-3′; EEA1 (N-term mNG11): crRNA sequence, 5′-GTGGTGGTTAAACCATGTTA-3′; donor oligonucleotide sequence, 5′-CCCGCGCAGGGTCTGGAGAGTCACCGCGGCGGCGCCGGGTGGTGGTTAAACCATGACCGAGCTCAACTTCAAGGAGTGGCAAAAGGCCTTTACCGATATGATGGGTGGCGGATTGGAAGTTTTGTTTCAAGGTCCAGGAAGTGGTTTAAGGAGGATTTTACAGAGGGTAAGAGAGTGAAACCGTCTTTCTGGCAGCACGT-3′

To generate CRISPRi- HEK293T cells that monitor endolysosomal damage, CRISPRi-HEK293T cells were transduced with the fluorescent Gal3 reporter (pJC41),^41^ and a polyclonal population was sorted by FACS.

The GFP-LC3-RFP-LC3∆G reporter line was generated as follows: HEK293T cells stably expressing dCas9-BFP-KRAB was transduced with pRT48 and a monoclonal line was selected for a high dynamic range for autophagic flux and high CRISPRi activity.

### Primary CRISPRi genome-wide screen

The genome-wide screen was carried out as seven parallel screens with sublibraries H1-H7.^40^ For each sublibrary, 100 million HEK293T cells were seeded into a 15 cm^2^ plate with complete DMEM and transfected with 5 μg of lentiviral plasmid packaging mix and 5 μg of pooled sgRNA library plasmid using lipofectamine 2000 (ThermoFisher Cat# 11668019) and incubated for 2 days.

Lentivirus-containing media was removed and filter sterilized using a Millex-GV syringe filter unit (Millipore Cat# SLGV033RB). For each library, 80 million FRET reporter cells were infected with filtered lentivirus-containing media for a target MOI of ~30-70%. Infected cells were selected with puromycin (1 ng/μL) for three days until % BFP >95%. Puromycin selected cells were then seeded at 25,000 cells/ well into fifty 96-well plates (ThermoFisher Cat# 12556008) with 0.25 ug/mL fibrils and 0.002% (v/v) lipofectamine. After 24 hours, cells were removed from the plates and sorted into FRET negative and FRET positive populations. Genomic DNA was isolated using the Macherey-Nagel Blood L kit (Machery-Nagel Cat# 740954.20) according to the manufacturer’s protocol. SgRNA-encoding cassettes were then amplified by PCR, purified using SPRI-select beads, and deep-sequenced as previously published.^23^ Phenotypes and P values for each gene were generated using the following pipeline: (https://kampmannlab.ucsf.edu/mageck-inc).^23^

### Microscopy and Flow cytometry secondary assays

To monitor α-synuclein aggregation, FRET reporter cells were seeded (25,000 cells/well) into 70 μL per well in a 96 well plates. In each well, 3 μL 0.25 mg/mL fibrils were complexed with 0.2 μL Lipofectamine 2000 for 1 hour at room temperature and added to 27 μL Optimem (ThermoFisher Cat# 31985070). The complex in Optimem was then added to pre-seeded reporter cells in 96-well plates and analyzed 24 hours after seeding. For analysis, cells were stained with Hoechst 33342 (ThermoFisher Cat# 5553141) at 0.5 μg/mL and analyzed by flow cytometry using a BD FACSCelesta or by fluorescence microscopy using an InCell 6000 (GE Cat# 28-9938-51). Cells with sgRNA knockdown were similarly analyzed using a comparable protocol 5 days after transduction with individual sgRNA-encoding lentivirus.

To monitor fibril uptake, CRISPRi-HEK293T cells were seeded (25,000 cells/well) into 70 μL in a 96-well plate with 3 μL of 0.25 mg/mL fibrils complexed with 0.2 μL Lipofectamine 2000 in 30 μL of Optimem, with various apilimod concentrations. Cells collected were collected at various timepoints until t = 6 hours after seeding for analysis by flow cytometry. Median mClover2 values were calculated in FlowJo and averaged between 3 technical replicates.

To monitor plasmid uptake, CRISPRi-HEK293T cells were seeded into 24-well plates at 200,000 cells/well and cultured for 24 hours. 5 ng of Plg15-BFP^18^ DNA complexed with 3 μL of Lipofetamine in 100 μL Optimem and various concentrations of apilimod was added to each well. After 24 hours cells were collected and analyzed via flow cytometry. Median BFP values were calculated in FlowJo and averaged between 2 technical replicates.

To monitor α-synuclein-Clover2 steady-state levels, FRET reporter was treated with various apilimod concentrations were seeded (25,000 cells/well) into 70 μL in a 96-well plate with 3 μL of 0.25 mg/mL fibrils was complexed with 0.2 μL Lipofectamine 2000 in 30 μL of Optimem. Median Clover2 values were calculated in FlowJo and averaged between 3 technical replicates.

To monitor autophagy, the CRISPRi-HEK293T cells expressing GFP-LC3-RFP-LC3∆G reporter were seeded (25,000 cells/well) into 70 μL in a 96-well plate with 3 μL of 0.25 mg/mL fibrils complexed with 0.2 μL Lipofectamine 2000 in 30 μL of Optimem, with various apilimod, bafilomycin and torin concentrations. After 24 hours cells were collected and analyzed via flow cytometry. The ratio of RFP/GFP was calculated in FlowJo and averaged between 3 technical replicates.

To monitor Gal3-EGFP puncta formation, CRISPRi-HEK293T cells expressing EGFP-Gal3 were seeded (25,000 cells/well) into 70 μL/ well in a 96-well black bottom plates, exposed to 3 μL of 0.25 mg/mL fibrils was complexed with 0.2 μL Lipofectamine 2000 in 30 μL of Optimem, and treated with LLOME or Lipofectamine at varying concentrations. 24 hours after seeding, cells were stained with Draq5 (Abcam, Cat# 108410) and images were collected by an InCell 6000. Images were analyzed using CellProfiler by quantifying the integrated density of identified Gal3 puncta divided by Draq5-stained nuclei and averaged between 3 images.

To track fibrils through the endolysosomal system in cells, LAMP1 and EEA1 were first pre-stained HEK293T cells with Hoechst diluted 1:10,000 in DMEM for 30 minutes at 37 °C, and first seeded (28,000 cells/well) into 150 μL microscopy media (DMEM, 10% FBS, 1% L-glutamine, 1% penicillin streptomycin, no phenol red) in a glass-bottom 96 well microscopy plate (Grenier, Cat# 655891) coated with fibronectin. The cells were then allowed to rest for half an hour at 37 °C. At the half hour mark, the cells were treated with either 360 nM apilimod or DMSO. The plate was then transferred to the microscope, and the pre-fibril images were acquired. Forty minutes after drug treatment, 30 uL of the fibril mixture (4 μL of 0.25 mg/mL Alexa 555-tagged fibrils complexed with 0.2 μL Lipofectamine 2000 in 28 μL of Optimem) was added to the cells. Z-stacks were then taken 30 minutes, 1 hour, 2 hours, 4 hours and 24 hours after the addition of fibrils using an Andor Dragonfly High Speed Confocal Platform. All images were acquired at 63× magnification.

### Filter-trap and immunoblot

FRET reporter cells were seeded (25,000 cells/well) in 70 μL per well into 96 well plates with various concentrations of apilimod. In each well 3 μL 0.25 mg/mL α-synuclein fibrils was complexed with 0.2 μL Lipofectamine 2000 for 1 hr at room temperature and added to 27 μL Optimem (ThermoFisher Cat# 31985070). Cells were collected by centrifugation at 1000 × *g* for 5 min and washed twice with PBS. Pellets were resuspended in 100 μL RIPA buffer containing protease inhibitors and homogenized on a tip sonicator (Sonopuls 2070) for 1 s at a power level of 10% for 5 s. 50 μL of sample was removed added to 50 μL 1× TBS or 1× TBS + 8M Urea mixed well. A dot blot vacuum manifold (Schleicher and Schuell Dot-Blot System) was used to pull 50 μl of each sample through onto a 0.45 μm nitrocellulose membrane (Biorad, Cat#162-0115). The membrane was washed 1× with water, blocked with Intercept Blocking Buffer (LiCor, Cat# 927-70001). Primary antibodies against human α-synuclein (Thermo Fisher, Cat# AHB0261) and GFP (Genetex, Cat# GTX113617) were used to detect proteins. Blots were then incubated with secondary antibodies (LiCor catalog numbers 926-32213 and 926-68072) and imaged on the Odyssey Fc Imaging system (LiCor, Cat# 2800). Immunoblot intensity was normalized to protein concentration as determined by a BCA Protein Assay Kit (Pierce, Cat# 23225). Images were processed and analyzed using LiCor Image Studio™ software.

### qRT-PCR

CRISPRi-HEK293T cells expressing a nontargeting or targeting sgRNA were collected by centrifugation at 1000 × *g* for 5 min and washed twice with PBS. RNA was purified using a RNA purification kit (Zymo, Cat#R1051). 500 ng of total RNA from each sample were reverse transcribed to cDNA using the Bioline SensiFAST cDNA Synthesis kit (Bioline, Cat# BIO-65054). The cDNA was diluted 13-fold in ddH2O and 4 uL of this dilution was used for each quantitative real-time PCR (qPCR). qPCR reactions were set up using SensiFast 2× Mastermix (Bioline Cat# BIO-94020) and oligonucleotides targeting genes of interest (IDT) in triplicate and run on the Quantstudio6 according to manufacturer’s protocols. Reactions were normalized to *GAPDH* as an internal loading control. sgRNA activity is expressed as knockdown efficiency. qPCR results and primer sequences are provided in Table S2.

## AUTHOR CONTRIBUTIONS

S.K.S. and M.K. conceptualization; S.K.S., M.C., S.B., R.T., A.W., E.T., I.E.J., C.N. E.N.M., J.S., M.D.L. and M.K. resources; S.K.S and M.K. formal analysis; S.K.S., M.C., S.B., R.T., A.W., E.T., I.E.J., C.N. E.N.M., J.S., M.D.L. and M.K. investigation; S.K.S and M.K. methodology; S.K.S and M.K. writing-original draft; S.K.S., M.C., S.B., R.T., A.W., E.T., I.E.J., C.N. E.N.M., J.S., D.R.S., J.E.S., M.D.L. and M.K. writing review and editing; R.T. and M.K. software; D.R.S., J.E.G., and M.K. funding acquisition; M.K. supervision.

## ACKNOWLEDGEMENTS

We thank Ana Maria Cuervo, Robert Edwards, Matt Jacobson, Carmen Nussbaum, Xiaoyan Guo, Nina Dräger, Emmy Li, Kun Leng, and other members of the Kampmann Lab for helpful discussions on experimental design and interpretation. We thank all co-authors, Avi Samelson, and Xiaoyan Guo for discussions and feedback on the manuscript. We thank Eric Chow and Derek Bogdanoff (UCSF Center for Advanced Technology) for support with next-generation sequencing and Sarah Elmes (UCSF Laboratory for Cell Analysis) for support with FACS. This work was supported by a Department of Defense NDSEG fellowship (S.K.S.), NIH grants DP2 GM119139 and R01 AG062359 (M.K.).

---

**Fig. S1.**
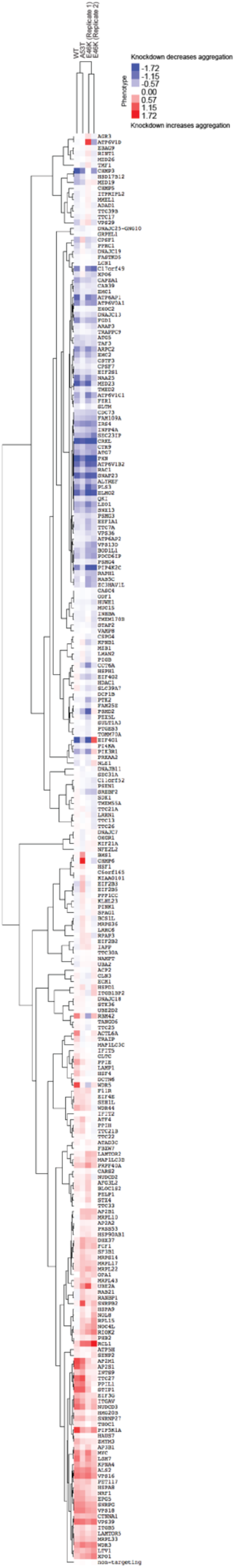
Full heat map displaying gene names and clustering analysis shown in Fig. 3A.

**Fig. S2.**
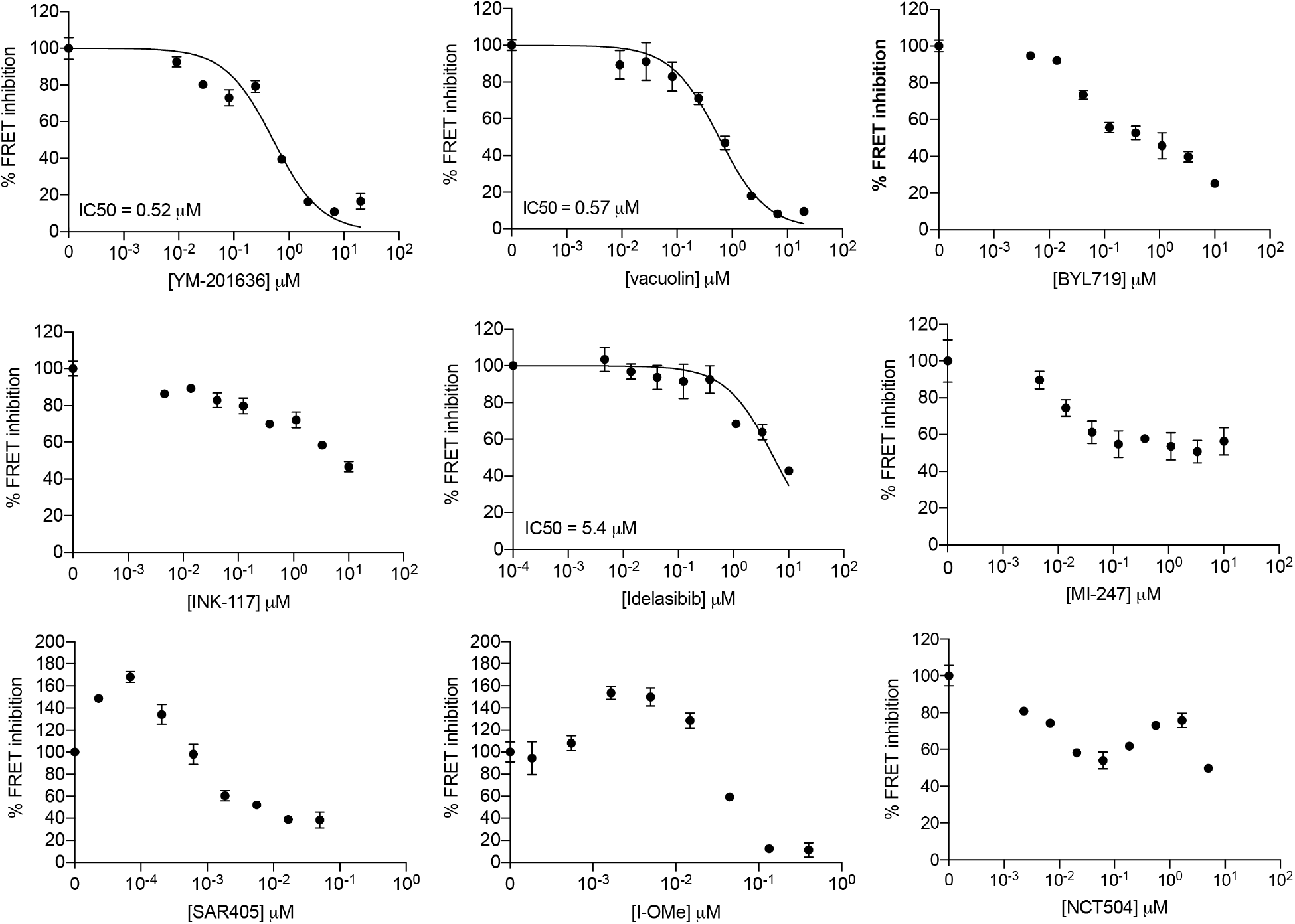
Aggregation efficiency in FRET reporter cells treated with various concentration of inhibitors of phosphatidylinositol signaling pathway members. Error bars represent mean ± s.e.m from n = 3 technical replicates.

## SUPPLEMENTAL TABLE LEGENDS

**Table S1: Phenotypes and p-values from the primary CRISPRi genome-wide screen for genes controlling α-synuclein aggregation.**

Whole Genome tab: Results from CRISPRi screens for were analyzed by the MAGeCK-iNC pipeline (see Methods for details) and are listed for all genes in the genome. In order to compare phenotypes between sublibraries screened on different days, phenotypes for each sgRNA within each sublibrary were normalized to the standard deviation of non-targeting control within each screen Columns are: Targeted transcription start site, knockdown phenotype (epsilon), P value, and Gene score.

Hits tab: Hit genes determined using an FDR<0.05 and their categorization for visualization in **Fig. 1B**. Columns are: Targeted transcription start site, knockdown phenotype (epsilon), P value, and Gene score, Gene categorization.

**Table S2: sgRNA protospacer sequences and phenotypes from focused CRISPRi retest screens, individually cloned sgRNA, their knockdown efficiency by qPCR, and qPCR primer sequences.**

Retest sgRNA tab: Protospacer sequences of the focused retest library shown in **Fig. 2**.

Retest clustering results tab: Relative phenotypes from focused screen in **Fig 2B**.

sgRNA oligos and qPCR tab: Oligonucleotide sequences used to clone sgRNAs and their knockdown efficiencies by qPCR.

qPCR primers tab: Oligonucleotide sequences used for qPCR experiments.

